# The genome of the colonial hydroid *Hydractinia* reveals their stem cells utilize a toolkit of evolutionarily shared genes with all animals

**DOI:** 10.1101/2023.08.25.554815

**Authors:** Christine E. Schnitzler, E. Sally Chang, Justin Waletich, Gonzalo Quiroga-Artigas, Wai Yee Wong, Anh-Dao Nguyen, Sofia N. Barreira, Liam Doonan, Paul Gonzalez, Sergey Koren, James M. Gahan, Steven M. Sanders, Brian Bradshaw, Timothy Q. DuBuc, Febrimarsa, Danielle de Jong, Eric P. Nawrocki, Alexandra Larson, Samantha Klasfeld, Sebastian G. Gornik, R. Travis Moreland, Tyra G. Wolfsberg, Adam M. Phillippy, James C. Mullikin, Oleg Simakov, Paulyn Cartwright, Matthew Nicotra, Uri Frank, Andreas D. Baxevanis

## Abstract

*Hydractinia* is a colonial marine hydroid that exhibits remarkable biological properties, including the capacity to regenerate its entire body throughout its lifetime, a process made possible by its adult migratory stem cells, known as i-cells. Here, we provide an in-depth characterization of the genomic structure and gene content of two *Hydractinia* species, *H. symbiolongicarpus* and *H. echinata*, placing them in a comparative evolutionary framework with other cnidarian genomes. We also generated and annotated a single-cell transcriptomic atlas for adult male *H. symbiolongicarpus* and identified cell type markers for all major cell types, including key i-cell markers. Orthology analyses based on the markers revealed that *Hydractinia*’s i-cells are highly enriched in genes that are widely shared amongst animals, a striking finding given that *Hydractinia* has a higher proportion of phylum-specific genes than any of the other 41 animals in our orthology analysis. These results indicate that *Hydractinia*’s stem cells and early progenitor cells may use a toolkit shared with all animals, making it a promising model organism for future exploration of stem cell biology and regenerative medicine. The genomic and transcriptomic resources for *Hydractinia* presented here will enable further studies of their regenerative capacity, colonial morphology, and ability to distinguish self from non-self.

## INTRODUCTION

*Hydractinia* is a small colonial marine invertebrate belonging to the phylum Cnidaria that grows on snail shells inhabited by hermit crabs, with polyps feeding opportunistically on small plankton and sharing resources throughout the colony. The polyp types found within these gonochoristic colonies include feeding polyps (gastrozooids), sexual polyps (gonozooids), and defensive polyps (dactylozooids and tentaculozooids). The colonies lend themselves to experimental study as they are easily cultured on glass microscope slides (Figure 1A). Marine hydroids, including *Hydractinia*, have fascinated biologists since at least the 1800s (Weismann 1883) due to their pluripotent adult stem cells (‘i-cells’) (Varley et al. 2023) and impressive regenerative capabilities. In fact, the term ‘stem cell’ (*stammzellen*) was coined by August Weismann in an 1883 chapter on *Hydractinia*’s putative migratory sperm progenitors (Weismann 1883; Wessel 2013). Other characteristics of these organisms such as allorecognition – a colony’s ability to distinguish itself from conspecifics – have also received considerable attention (Nicotra 2019). Their closest well-studied relative is the freshwater *Hydra*, which shares many characteristics with *Hydractinia*, including i-cells, the capacity for whole-body regeneration, and the absence of a medusa adult phase. However, *Hydractinia* differs from *Hydra* in several important respects, including its colonial morphology, polyp polymorphism, and possession of a single self-renewing stem cell lineage (Varley et al. 2023) as compared to the three self-renewing lineages in *Hydra* (interstitial, endodermal, and ectodermal). There are also salient differences in their life cycles, with *Hydractinia* undergoing metamorphosis from the larval to adult form, whereas *Hydra* exhibits direct development with no larval stage. These differences between the two lineages are unsurprising given that they diverged at least 500 MYA (Steele et al. 2011).

**Figure 1:**
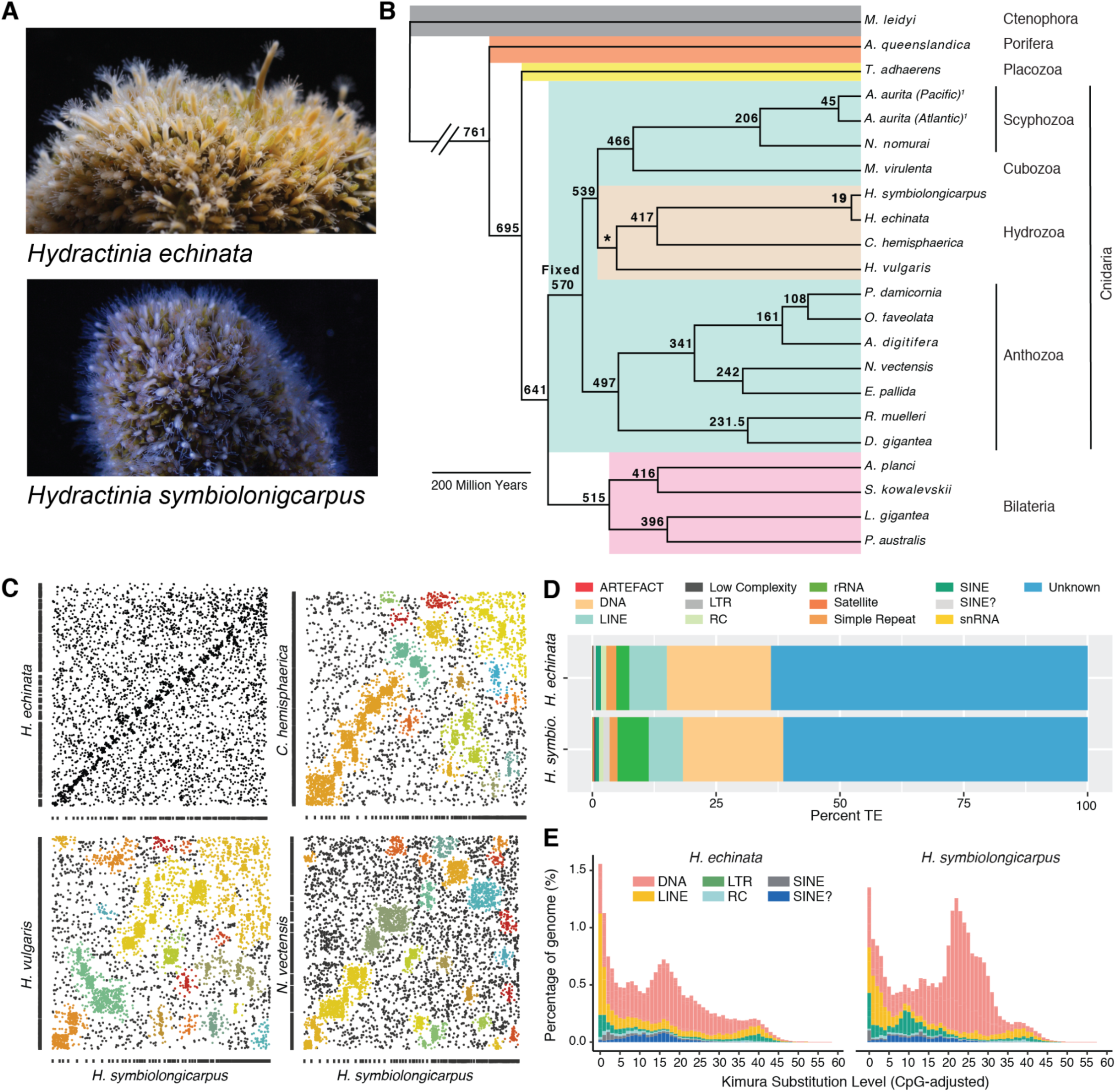
Overview of *Hydractinia*, phylogenetic analysis, synteny analysis, and analysis of repetitive elements. (A) *Hydractinia echinata* colony (top panel); *Hydractinia symbiolongicarpus* colony (bottom panel). (B) Maximum likelihood phylogeny estimated from a data set of single copy orthologs as inferred by Orthofinder2 showing that the two *Hydractinia* species cluster together with *C. hemisphaerica* and *H. vulgaris* branching next to them within the Hydrozoa. Divergence times were estimated using the r8s program. The age of Cnidaria was fixed at 570 MYA and the age of Hydrozoa constrained to 500 MYA based upon Cartwright and Collins (2007). (C) Syntenic dot plots comparing *H. symbiolongicarpus* with four cnidarian species: *H. echinata*; *C. hemisphaerica; H. vulgaris*; and *N.vectensis*. Colored boxes indicate linkage groups. (D) Stacked bar chart showing proportions of different transposable elements classes in each *Hydractinia* genome using RepeatMasker de novo analysis. (E) Repeat landscape analysis showing overall a highly similar evolutionary history of invasion of repetitive elements in the two species. In *H. symbiolongicarpus*, there was a species-specific recent expansion (at approximately 10% nucleotide substitution) of LTR retrotransposons.

Here, we report highly contiguous genome assemblies for two species: *H. symbiolongicarpus*, found along the east coast of the United States, and *H. echinata*, found in European waters, comparing their genome structure and content with those of other cnidarians and other animals. We performed orthology inference analyses using the predicted proteomes of the two *Hydractinia* species, along with proteomes from 41 additional animal species and six related eukaryotes.

These data were then used to describe overall gene evolutionary patterns, including lineage specificity and gene family dynamics. Phylogenetic analyses based on conserved single-copy genes agreed with previous placements of *Hydractinia* within the hydrozoan cnidarians, positioning *Hydractinia* together with *Clytia hemisphaerica* and *Hydra vulgaris*. In addition, divergence time analysis yielded estimates of when the two *Hydractinia* species diverged from other cnidarians. We also analyzed gene synteny between the two species and with other cnidarian genomes and characterized the repeat content and diversity of the two species. We explored *Hydractinia*’s mitochondrial genome structure, comparing it to those of other cnidarian species, and deduced the presence and absence of homeodomain-containing genes in these species.

We used a single-cell transcriptomic approach to create a robust cell-type atlas that allowed us to annotate all clusters and identify specific genes that define individual cell types in adult animals. We further explored our cell type marker lists using orthology assignments to identify cell type markers that are evolutionarily conserved with other animals. We found that i-cells and progenitors are defined by genes that are highly conserved among animals, in contrast to most other cell types that contain a significant proportion of cnidarian-specific genes. Together, these data mark a new era in the exploration of this remarkable marine hydrozoan, providing new insights into the diversity of cnidarian genomes and our view of their evolution.

## RESULTS

### Sequencing, assembly, and annotation of Hydractinia genomes

We estimated the genome sizes for *Hydractinia symbiolongicarpus* male wildtype strain 291-10, *Hydractinia echinata* female wildtype strain F4, the closely related hydrozoan *Podocoryna carnea* male wildtype strain PcLH01, and *Hydra vulgaris* strain 105 using propidium iodide staining of isolated nuclei followed by flow cytometric analysis (Hare and Johnston 2011) (detailed methods in Supplement). The resulting genome size estimates were 514 Mb for *H. symbiolongicarpus* and 775 Mb for *H. echinata* (Table S1). By way of comparison, our estimate for *P. carnea* was 517 Mb and 1086 Mb for *H. vulgaris*, consistent with previous reports (Chapman et al. 2010). We then isolated high molecular weight DNA from adult polyps and sequenced both *Hydractinia* genomes using a combination of PacBio SMRT long-read and Illumina short-read sequence data (detailed methods in Supplement). The PacBio libraries had insert sizes ranging from 6.9-10.6 kb and each library was sequenced across several SMRT cells (Table S2). These PacBio data were then used to generate primary contig assemblies using the diploid-aware assembler Canu (Koren et al. 2017) (Table S3).

Assemblies were also generated with Falcon_unzip (Chin et al. 2016) but these were ultimately abandoned after comparison with the Canu assemblies (Table S4). Canu attempts to assemble and phase contigs representing alternative haplotypes in heterozygous regions into primary and secondary assemblies via a filtering step, but this phasing can be challenging when applied to genomes that exhibit a high level of heterozygosity. Here, we estimated overall heterozygosity to be 1.33% for *H. symbiolongicarpus* and 0.85% for *H. echinata* (Figure S1). In addition, Canu phasing resulted in primary assemblies that had many duplicated loci, with initial BUSCO (Simão et al. 2015) analyses indicating 42% and 29% duplicated genes in the *H. symbiolongicarpus* and *H. echinata* assemblies, respectively. To address this, we used MUMmer 3.23 (Kurtz et al. 2004) to better-separate haplotypes (detailed methods in Supplement). Following this contig filtering procedure, the presence of duplicated loci in the primary assemblies was reduced to 11% for *H. symbiolongicarpus* and 10% for *H. echinata*. These primary contig assemblies were then scaffolded with Illumina Chicago libraries through Dovetail HiRise scaffolding (Putnam et al. 2016), then gap-filled using PBJelly (English et al. 2012). The assemblies were polished using the final consensus-calling algorithm Arrow (Chin et al. 2013) and further polished with Pilon (Walker et al. 2014). The resulting final scaffolded and polished primary assemblies resulted in a 406 Mb assembly for *H. symbiolongicarpus* consisting of 4,840 scaffolds with a scaffold N50 of 2,236 kb, and a 565 Mb assembly for *H. echinata* consisting of 7,767 scaffolds with a scaffold N50 of 904 kb (Table S3). BUSCO percentages for the final assemblies indicated a high level of completeness for both genomes (89.6% for *H. symbiolongicarpus* and 89.1% for *H. echinata*; Table S3). Karyotype analysis of *H. symbiolongicarpus* previously reported 15 pairs of chromosomes (2n = 30) for this species (Chen et al. 2023), consistent with the chromosome count of several other cnidarians, including *H. vulgaris, C. hemisphaerica* (Munro et al. 2023) and *Nematostella vectensis* (Zacharias et al. 2004; Putnam et al. 2007; Guo et al. 2018).

### Gene model prediction and annotation

Using RNA-seq reads and assembled transcripts from adult animals to guide the annotation process, we predicted genes for each genome using Augustus (Haas et al. 2008), with detailed methods provided in Files S1 and S2 and summary statistics in File S3. 22,022 genes were predicted for *H. symbiolongicarpus* and 28,825 for *H. echinata*. Coding regions make up about 8% of each assembly, while noncoding regions account for 92%. On average, *H. symbiolongicarpus* has 7.47 exons and 6.47 exons per gene, compared to 6.60 exons and 5.60 introns per gene in *H. echinata* (Table S5). The average intergenic region is 6,679 bp for *H. symbiolongicarpus* and 7,603 bp for *H. echinata* (Table S5). 5’ and 3’ UTR predictions were performed with PASA (Haas et al. 2008), indicating that 48% (*H. symbiolongicarpus*) and 42% (*H. echinata*) of the gene models have predicted UTRs. Some *Hydractinia* transcripts undergo trans-spliced leader addition processing that is known to occur in hydrozoan genomes (Stover and Steele 2001; Derelle et al. 2010). The replacement of 5’ UTR sequences by short sequences that are trans-spliced from non-coding spliced leader RNAs occurs in a few distantly related animal groups (Hastings 2005). We detected spliced leader sequences in our mRNA sequencing data, as well as spliced leader genes. Our ability to accurately predict 5’ UTRs for some gene models was likely impacted by this phenomenon.

We evaluated completeness of the predicted gene models via BUSCO v5 (*9*) with the Metazoa dataset of 954 proteins. For *H. symbiolongicarpus*, there were 92.5% complete and 10.2% duplicated genes (Tables S3 and S12 tab SM1), while there were 90.7% complete and 12.3% duplicated genes in *H. echinata* (Tables S3 and S12 tab SM1). We determined the percentage of gene models that had assembled transcript support and performed functional annotation on these gene models, combining our RNA-seq data from adult animals with additional RNA-seq data from *H. symbiolongicarpus* developmental stages or *H. echinata* polyp head regeneration timepoints (details in Supplement, Files S4-S6, Tables S6-S7) for our transcript support analysis. Overall, 78% of *H. symbiolongicarpus* gene models and 63% of *H. echinata* gene models had transcript support with at least 90% gene overlap (Figures S2-5, Table S8). A small percentage of gene models had no overlapping transcript support (14% *H. symbiolongicarpus*, 21.5% *H. echinata*; Figures S2 and S4, Table S8). Functional annotation of gene models was performed using several approaches that included a DIAMOND search (Buchfink et al. 2015) of NCBI’s nr database and using PANNZER2 (Törönen et al. 2018) (Table S9). Additional details on the annotation process are provided in the supplement. Overall, 88.5% of *H. symbiolongicarpus* gene models and 76.2% of *H. echinata* gene models had some level of annotation: a DIAMOND hit to NCBI nr, a PANNZER2 hit, or both (Table S9).

### Mitochondrial Genome

Cnidarians are characterized by mitochondrial genomic diversity, varying in overall mtDNA conformation (circular or linear), gene content, gene organization, and the number of mitochondrial chromosomes within each species (Kayal et al. 2012; Smith et al. 2012; Kayal et al. 2015a). Medusozoan cnidarians possess linear monomeric or multimeric mitochondrial chromosomes, while most anthozoan cnidarians possess circular mtDNA (Figure S6) (Kayal et al. 2012, 2015b; Bridge et al. 1992; Brugler and France 2007). The typical mtDNA observed in cnidarians consists of a set of 17 genes: the small and large ribosomal genes, methionine and tryptophan transfer RNA genes, and 13 energy pathway proteins (Bridge et al. 1992; Beagley et al. 1998). These genes are usually organized in the same transcriptional orientation, with a partial or complete extra copy of the *Cox1* gene in the opposite transcriptional orientation at one end of the chromosome (Kayal and Lavrov 2008). Secondary structures in intergenic regions and at the ends of the mtDNA regions may be involved in the control of replication and transcription (Brugler and France 2007; Stampar et al. 2019) and are also thought to protect the ends of the mitochondrial chromosome given their lack of traditional telomeric repeats, as previously observed in *Hydra oligactis* (Kayal et al. 2012; Smith et al. 2012; Brugler and France 2007; Beagley et al. 1998). Furthermore, introns, duplicated genes, and several additional protein-coding genes have been observed in several non-hydrozoan cnidarian mitogenomes (Beagley et al. 1998; Shao et al. 2006; Chen et al. 2008; Voigt et al. 2008).

The linear mitochondrial genome of *Hydractinia* is located on a single scaffold in both *Hydractinia* species, containing the coding sequences for the large (*16S/RNL*) and small (*12S/RNL*) ribosomal subunits, mitochondrial transfer RNA (tRNA) genes, all cnidarian mitochondrial proteins (*Cox1-3*, *Cob*, *Nad1-6*, and *Nad4L*), and inverted terminal repeats (ITRs) that form G-rich loops at both ends of the molecule. This strongly suggests that *Hydractinia* contains only one mitochondrial chromosome, similar to what has been observed in other hydrozoan genomes (Figure S7, Table S10) (Kayal et al. 2012; Smith et al. 2012; Kayal and Lavrov 2008). *Hydractinia*’s mitochondria are mostly devoid of tRNAs, with both species containing just one tRNA-Met sequence and one tRNA-Trp sequence (Figure S8). These sequences form the characteristic tRNA hairpin structure and are in non-coding regions (Figure S8). An alternative mechanism for the replication and expression of linear mitochondrial genomes has been suggested, where transcription and replication occur in two directions, starting from a large intergenic spacer (Kayal et al. 2015a). The origin of replication (Ori) is characterized by stable stem-loop configurations containing T-rich loops and abrupt changes in DNA composition bias (Brugler and France 2007; Stampar et al. 2019). Based on these characteristics, we propose that the origin of replication in *Hydractinia* is in the intergenic spacer between the large ribosomal subunit (*16S/RNL*) and the *Cox2* gene (Figures S7 and S9). The ITRs of both *Hydractinia* species can form G-rich loops that likely protect the ends of these linear mitochondrial chromosomes in the absence of telomeric sequences (Figure S10). In addition, the presence of non-functional (and gradually degrading) nuclear copies of mtDNA (NUMTs) have previously been identified in *H. vulgaris* (Song et al. 2013). Sequence similarity searches did not detect NUMTs within either *Hydractinia* genome. This result was confirmed by the lack of sequence variance in Illumina raw reads mapped to their mitochondrial genomes. Other cnidarians with linear mtDNAs, such as the jellyfish *Sanderia malayensis* and *Rhopilema esculentum*, were also shown to not contain NUMTs (Nong et al. 2020).

### Orthology inference, phylogenetic analyses, and divergence time estimates

Orthology inference analysis was performed on a splice-filtered dataset (File S7) consisting of proteomes from 49 eukaryotic species encompassing 15 animal phyla and four non-animal outgroups (detailed methods in Supplement; Table S11). Taxon selection was initially based on a data set used by Maxwell et al. (Maxwell et al. 2014) to infer the evolutionary origins of human disease-associated gene families that was then expanded to place the *Hydractinia* genomes in an evolutionary context with other cnidarian genomes. To that end, 16 cnidarian species spread across the main cnidarian lineages were included. This represents the largest sampling of cnidarians in any genome-wide orthology inference study performed to date and provides increased resolution for characterizing evolutionary dynamics among cnidarians, as well as between cnidarians and other animals. An input species tree (Figure S11) based on the current literature was provided to OrthoFinder2. A total of 33,325 orthogroups containing 81.2% of the proteins were recovered in the dataset. These orthogroups were then used as the basis for the analyses described below (Files S8-S14).

For our phylogenetic analysis, we selected a subset of single copy ortholog (SCO) sequences from our orthogroup data set (File S15). These SCOs were chosen for their presence in at least 12 of 15 cnidarian species; four bilaterian and three non-bilaterian outgroup species that also contained these SCOs were included in the analysis. The final concatenated, aligned, and trimmed data set included sequences from 216 orthogroups, resulting in an alignment of 50,457 nucleotides (File S16). The resulting maximum likelihood tree, generated using IQ-Tree2 (Files S17-S18), confirmed known relationships within Cnidaria, including placing the two *Hydractinia* species closest to *C. hemisphaerica* (Figure 1B). This tree was then used to estimate divergence times within the phylum using r8S (Sanderson 2003). Our age estimate for the most recent common ancestor of anthozoans is 496.6 MYA, while that of medusozoans is 538.9 MYA. Strikingly, while the estimated ages for clades within Cnidaria tend to be older than those previously reported (Khalturin et al. 2019), we find that the divergence time between the two *Hydractinia* species to be just 19.16 MYA (Figure 1B).

This estimate is much shorter than the estimated divergence times between lineages of the moon jelly *Aurelia aurita* [45.35 MYA in our study; 51-193 MYA reported by Khalturin et al. (Khalturin et al. 2019)] and is more comparable to the divergence time between lineages of *Hydra vulgaris* [10-16 MYA; (Wong et al. 2019)]. Providing an alternative input species tree with Porifera at the base did not significantly alter overall results of orthology inference or divergence time estimates (Figure S12).

### Synteny

We performed pairwise macrosynteny analyses comparing *H. symbiolongicarpus* and *H. echinata*, as well as a series of comparisons between each *Hydractinia* species and *C. hemisphaerica*, *H. vulgaris*, and *N. vectensis* by clustering scaffolds of these species based on the shared orthogroup numbers (detailed methods in Supplement and Files S19-S22). Despite not having chromosomal-level assemblies, we observed local collinearity between the two *Hydractinia* species (Figure 1C) and general chromosomal-level conservation beyond scaffold boundaries, as evidenced by scaffold clustering within the *Hydractinia* genus and beyond (Figure 1C). This indicates a high degree of synteny between the two *Hydractinia* species, an observation that is not surprising due to their close phylogenetic relationship and relatively recent divergence (Figure 1B). The observation that this conservation is shared with at least three other cnidarian species (Figure 1C, Figure S13) suggests that *Hydractinia* chromosomes show a similar degree of ancestrality (Simakov et al. 2022). Further chromosomal-level assembly and analysis will be required to validate this hypothesis and identify features unique to *Hydractinia*.

### Characterization of genomic repeats, including transposable elements

According to our RepeatMasker *de novo* analysis, genomic repeats comprise 55% of the *H. echinata* genome and 50% of the *H. symbiolongicarpus* genome. These figures are slightly lower than the percentage of repetitive DNA found in *H. vulgaris* (57%) but higher than that found in both *C. hemisphaerica* (39%) and *N. vectensis* (25%) [Table S12; (Chapman et al. 2010; Putnam et al. 2007; Leclère et al. 2019)]. The overall composition of repeat classes is similar between the two *Hydractinia* species (Figure 1D, Figure S14, Tables S13-S16). The largest proportion of repeats are unclassified in both genomes, accounting for around 60% of all repetitive elements; these unclassified repeats comprise 35% and 30% of the *H. echinata* and *H. symbiolongicarpus* genomes, respectively.

Beyond the unclassified repeats, DNA transposons comprise the most abundant class of transposable element, accounting for about 20% of all repetitive elements and 11% of each genome. This is similar to what has been observed in both *N. vectensis* and *H. vulgaris*, where DNA transposons are the most abundant class of transposable elements.

There were some dramatic differences in several DNA transposon superfamilies between the two species (Figure S14). Long interspersed nuclear elements (LINEs) accounted for 7% of all repetitive elements and 4% of each genome. Other repetitive element classes have similar compositions in the two genomes, except for long terminal repeat (LTR) retrotransposons. Although LTR retrotransposons only accounted for a small fraction of the genome in both species, there are some significant differences in their family composition and evolution between the species (Figure S14). The LTR retrotransposons accounted for 2.6% of all repetitive elements in *H. echinata* and 3% in *H. symbiolongicarpus*, representing 1.5% and 3% of these genomes, respectively. We performed a repeat landscape analysis (detailed methods in Supplement) that suggests a highly similar evolutionary history of invasion of repetitive elements in the two species (Figure 1E, Files S23-S24) with a few notable differences (Figures S15-S16). In *H. symbiolongicarpus*, there was a species-specific recent expansion (at ∼10% nucleotide substitution) of LTR retrotransposons (Figure S16). This small expansion was mainly composed of members of the Gypsy family of LTRs. The two genomes also harbor different types of endogenous retroviruses (ERVs). Endogenous retrovirus group K genes (ERKVs) are only present in *H. echinata*, whereas endogenous retrovirus group L genes (ERVLs) are only present in *H. symbiolongicarpus*, suggesting two recent independent invasions of ERVs after the speciation event around 19 MYA (Figure S16).

### Orthogroup lineage specificity and overall patterns of evolutionary novelty

Recent cnidarian genome sequencing projects (Khalturin et al. 2019; Leclère et al. 2019; Gold et al. 2019) have demonstrated the contribution of both taxon-restricted and shared ancestral gene families to cnidarian-specific cell types, such as those found in the medusa. To evaluate the contribution of such gene families to evolutionary novelty in *Hydractinia*, we identified lineage-specific subsets of orthogroups. Out of the 33,325 orthogroups inferred by Orthofinder, roughly 26% are cnidarian-specific, 16% are medusozoan-specific, 8% are hydrozoan-specific, 6% are specific to *Hydractinia* + *Clytia*, and just under 5% are specific to the genus *Hydractinia*. In comparison, only 7% of orthogroups are specific to anthozoans. *H. echinata* possesses 46 species-specific orthogroups, while *H. symbiolongicarpus* possesses just 15 such orthogroups. Additionally, based on our sampling of 23 bilaterian species from a variety of phyla, the percentage of bilaterian-specific orthogroups (∼24%) is similar to the 26% found in cnidarians. This observation implies that the evolutionary novelty of orthogroups found in all of Bilateria is equal to that found just within the Cnidaria.

To evaluate the contribution of conserved gene families to *Hydractinia*’s evolution and further evaluate the broad suitability of cnidarians as animal models, we calculated the overlaps of orthogroups between major groups of cnidarians and bilaterians (Figure S17, Files S25-S27). At the broadest scale, cnidarians and bilaterians possess more shared than unshared orthogroups. This supports previous observations based on the genome sequences of *Hydra* (Chapman et al. 2010) and *Nematostella* (Putnam et al. 2007) that much of the cnidarian toolkit predates the divergence of Cnidaria and Bilateria. Splitting Cnidaria further into the Medusozoa and Anthozoa (Figure S17A), we observe that the number of orthogroups unique to Medusozoa + Bilateria is nearly equal to that for Anthozoa + Bilateria, both of which are greater than the number for Medusozoa + Anthozoa. This is consistent with numerous observations of deep divergence between medusozoan and anthozoan genomes, from fossil estimates to divergence time estimates (Steele et al. 2011; Khalturin et al. 2019).

To further investigate potential sources of evolutionary novelty, we calculated the percentage of genes within each species that are assigned to orthogroups that are species-specific, the percentage of phylum-specific and metazoan-specific genes, and the percentage of genes unassigned to an orthogroup. These five proportions are visualized in the right panel of Figure 2 for the 15 cnidarian species that were analyzed further using CAFE (see below). Proportions for all metazoan species in our analysis are visualized in Figure S18. Notably, *H. symbiolongicarpus* and *H. echinata* contain the highest percentages of phylum-specific genes of all 43 metazoan species we examined (23% and 22%, respectively), thereby indicating that their genomes contain the highest percentage of cnidarian-specific genes of all cnidarians included in this analysis. Coupled with the fact they possess relatively few species-specific orthogroups, this suggests that a significant proportion of their proteomes may have evolved at the genus, family, or subphylum level, which are grouped together under ‘Phylum-specific’ in the analysis featured in Figure 2. Additionally, a DIAMOND search indicated that most (90%) unassigned *Hydractinia* genes had no match in the NCBI nr database (Table S11 tab X.4). Transcript support for these genes (Table S8) indicates that a large proportion of these genes have >90% transcript overlap (51.28% in *H. symbiolongicarpus* and 35.35% in *H. echinata*) and are expressed by the animal. Thus, the two *Hydractinia* genomes appear to contain an abundance of evolutionarily novel genes.

**Figure 2:**
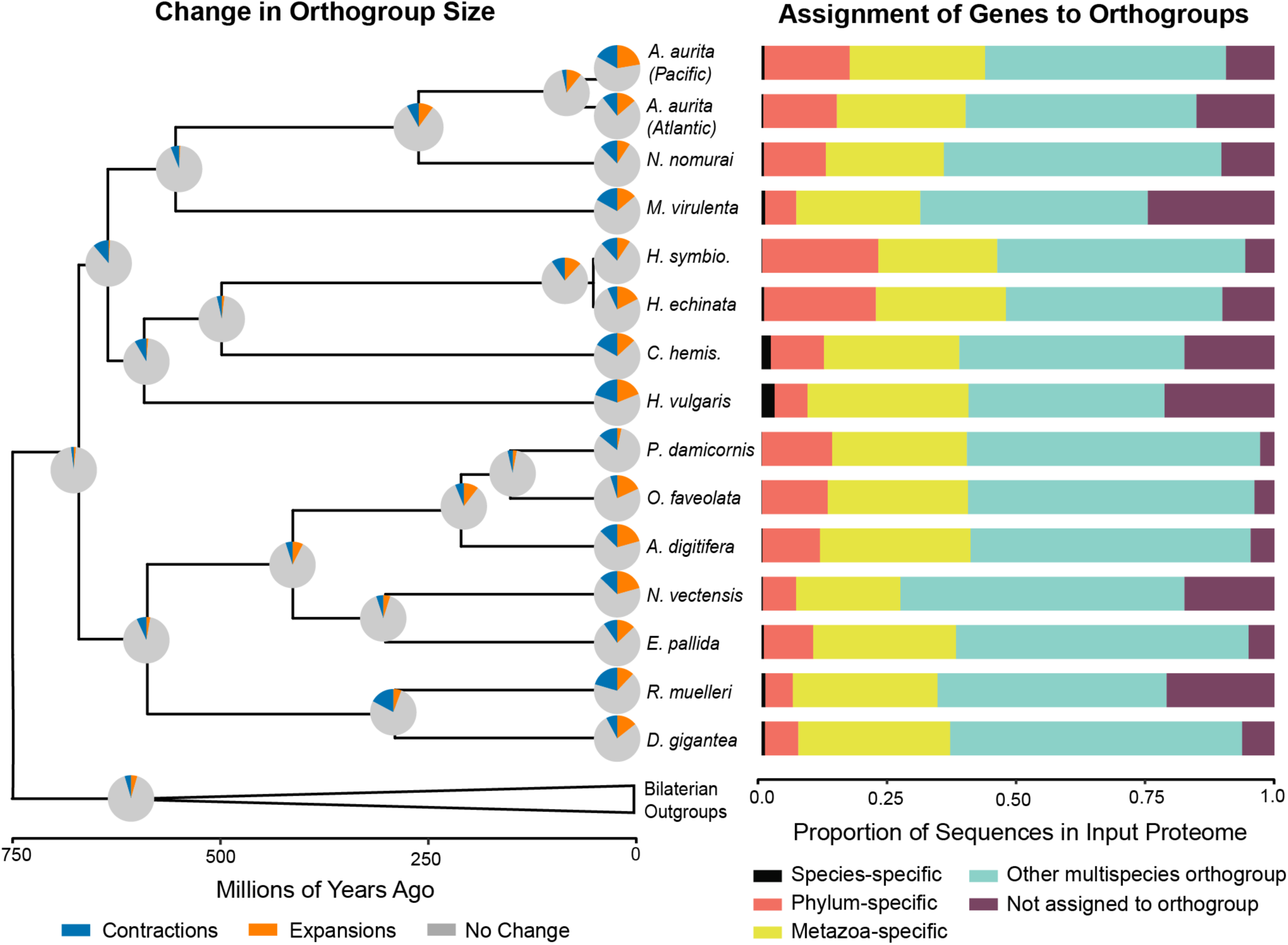
Summary of orthogroup evolution across a subset of sampled taxa. Left: Changes in gene family size estimated using CAFE. Pie charts represent changes along the branch leading to a given node or tip for all 8,433 orthogroups inferred to be present in the common ancestor of this tree. Branch lengths are as depicted in Figure 1B. Right: Proportion of input proteome sequences assigned by Orthofinder2 to different orthogroup categories. See Figure S16 for results for every species included in Orthofinder analysis and Table S12 tab SM1 for the number of input sequences in each proteome. The data used to create these figures can be found in Table S12 tabs X.2-3. ^1^Gold et al. 2019, ^2^Baltic/Atlantic genome from Khalturin et al. 2019.

### Estimating the evolutionary dynamics of gene families using CAFE

Focusing just on the Cnidaria + Bilateria subtree (19 species) derived from the 22 species tree inferred using IQ-Tree2 and r8s (described above), we estimated the evolutionary dynamics of the 8,433 Orthofinder-inferred orthogroups that are present in the ancestor of this subtree (additional details in Supplement, Files S28-S39). Using CAFE, gene family dynamics were estimated for each node and terminal taxon (De Bie et al. 2006; Han et al. 2013) in our subtree and are summarized as Figure 2 (left panel), with additional details available in Table S11, Tab X.8.

Across the whole tree (Figure 2), more changes in gene family size take place on the terminal branches of the tree than in the internal branches of the tree. Terminal branches have significantly more gene expansion or contraction as compared to internal branches [mean(terminal) = 2,375.7, mean(internal) = 1007, t = –8.5139, df = 33.99, p-value = 6.07 x 10^-10^]. This pattern is very clear when comparing the internal nodes of the cnidarian phylum with the terminal branches of this group (Figure 2). Of the 8,433 analyzed orthogroups, a total of 592 were found to be evolving rapidly on the subtree (Viterbi p-value <= 0.05). The distribution of these uniquely fast-evolving gene families per taxon/node can be found in Table S11, Tab X.8, and information about their putative identities can be found in the Supplement.

### Comparing evolutionary dynamics of H. symbiolongicarpus and H. echinata using CAFE

Roughly half of the orthogroups present in the *Hydractinia* genomes and included in the CAFE analysis have undergone some change in size (50% in *H. symbiolongicarpus* and 54% in *H. echinata*) when comparing their observed size to the inferred size of these orthogroups in the Cnidarian+Bilaterian ancestor. Notably, the two *Hydractinia* genomes have very different proportions of gains versus losses over their terminal branches. *H. echinata* has experienced more expansions with a higher number of genes per expansion, resulting in *H. echinata* gaining about twice as many (1.97x) individual gene copies in the past 19 million years. Conversely, *H. symbiolongicarpus* has a higher number of contracted gene families and has lost more genes per contraction, meaning that *H. symbiolongicarpus* has lost nearly 2.5 times more genes in total than *H. echinata* has since their divergence. Additionally, although *H. echinata* and *H. symbiolongicarpus* have lost 248 and 252 gene families, respectively, the identities of the lost families do not overlap at all. This implies that these species have undergone very different evolutionary trajectories since their divergence roughly 19 MYA. We performed additional comparisons of evolutionary dynamics in *Hydractinia* versus the other hydrozoan taxa (*H. vulgaris* and *C. hemisphaerica*) and versus the genus *Aurelia* (details in Supplement). Overall, *H. vulgaris* and *C. hemisphaerica* have more taxon-specific orthogroup size changes than either species of *Hydractinia*. However, when combining data from the two *Hydractinia* species to look at changes at the genus level, the number of changes are roughly similar between these hydrozoans. For the comparison with *Aurelia*, we found that the overall proportions of gains versus losses was much more similar between the two *Aurelia* lineages, in contrast with what we found for the two *Hydractinia* species (additional details in Supplement, Figure S19).

### The Non-coding RNA landscape: miRNAs

microRNAs (miRNAs) constitute a unique class of small non-coding RNAs of approximately 22 nucleotides (nt) in size that play crucial roles in development, cellular differentiation, and stress response in both plants and animals (Wheeler et al. 2009). Several studies have investigated miRNAs and the miRNA pathway in cnidarians (Moran et al. 2013; Praher et al. 2021). We generated small RNA-seq libraries for five samples of adult *H. echinata* polyps that were then sequenced (detailed methods in Supplement). The resulting reads were trimmed and mapped to the *H. echinata* genome using the miRDeep2 mapping algorithm (Friedländer et al. 2012). After mapping, the miRDeep2 algorithm predicted 347 miRNAs. Predictions with a score >5.0 were retained. To find the highest quality predicted miRNAs from this set, we performed custom automated filtering of the miRDeep2 output and then manually screened the filtered predictions (detailed methods in Supplement, Figure S20). 104 predictions passed our custom automated filtering. During manual screening, 48 of these were deemed to be high quality predictions, whereas 23 were found to be of low quality (Figure S21). After removing redundancy, we generated a final list of 38 unique high-quality mature miRNA sequences (Table S17). Of these, three are homologous to known cnidarian miRNAs (miR-2022, miR-2025, and miR-2030), with alignments shown in Figure S22. Figure S23 depicts a proposed evolutionary scenario for miRNAs in cnidarians that includes these new data from *H. echinata*.

### The Non-coding RNA landscape: rRNAs, tRNAs, and snoRNAs

In an attempt to provide the first detailed description of the non-coding RNA (ncRNA) landscape of any cnidarian species, we found that the two *Hydractinia* genomes encode the expected suite of functional non-coding RNAs commonly present in metazoan genomes. These included ribosomal RNA genes (rRNAs), transfer RNAs (tRNAs) for each amino acid isotype, spliceosomal RNAs for both the major (U1, U2, U4, U5 and U6) and minor spliceosome (U11, U12, U4atac, and U6atac), small nucleolar RNAs (snoRNAs), SRP RNA, RNase P RNA, RNase MRP RNA, and Vault RNA (Table S18). This characterization was based on results from Rfam (Kalvari et al. 2018), Infernal (Nawrocki and Eddy 2013) and tRNAscan-SE (Chan et al. 2021) as described in the Supplement. An unusual feature of many of these ncRNAs is their apparent organization into roughly evenly spaced tandem arrays of tens or even hundreds of nearly identical or highly similar copies. Each of these copies is separated by spacer regions ranging in length from several hundred to a few thousand nucleotides that are nearly identical or highly similar to one another (see Tables S19-S21 and Files S40-S48). In both *Hydractinia* genomes, these arrays include ribosomal RNAs, four of the five RNA components of the major spliceosome (U1, U2, U5, and U6), the small nucleolar RNA U3, and tRNAs for each of the 20 amino acids (Table S18, Tables S21-25). Although tandem arrays of some RNA genes – especially clusters of ribosomal RNA genes collectively known as rDNA – are common in eukaryotes (Cloix et al. 2000; Long and Dawid 1980), tandem array organization of tRNAs (Bermudez-Santana et al. 2010) is unusual outside of the *Entamoeba* genus of *Amoebozoa* (Tawari et al. 2008), with only one such example having been observed in mammals (Darrow and Chadwick 2014). The ncRNA tandem arrays only make up a small percentage of all regions that appear in tandem repeats in the *Hydractinia* assemblies. Tandem repeat regions detected using TRF (Benson 1999) having seven or more copies with a period length of 50 nt and ≥75% average similarity between repeats cover 18.7% of the *H. echinata* and 15.7% of the *H. symbiolongicarpus* assemblies. These TRF-defined repeats are largely a subset of the unclassified repeats identified by our RepeatMasker analysis detailed above (88.1% of the *H. echinata* and 72.0% of the *H. symbiolongicarpus* nucleotides in the TRF-defined repeat regions also exist in the unclassified repeat regions). The nucleotides covered by the RNA tandem arrays account for only 4.8% and 7.7% of these TRF-defined repetitive regions in *H. echinata* and *H. symbiolongicarpus*, respectively. While the biological significance of these ncRNA tandem arrays and other tandem repeat regions remains unclear in the absence of functional data, two important observations argue against the presence of these RNA tandem arrays being due to sequencing or assembly artifacts. First, when comparing these results to other cnidarian species, we were able to identify tandem arrays of 5S rRNA, tRNA, and U5 RNA in the *N. vectensis* genome (Putnam et al. 2007) but did not find RNA tandem arrays in other cnidarian genomes. Secondly, the draft genome assembly of *H. echinata*, sequenced and assembled using different methods (Török et al. 2016) than the primary *H. echinata* assembly presented here, also includes tandem arrays of 5S rRNA, SRP RNA, and tRNA and a significant fraction of that assembly is also in TRF-defined tandem repeats (5.1% of the genome). Taken together, this first characterization of the ncRNA landscape hints at interesting differences between cnidarians and other eukaryotes.

### The homeobox gene complement of Hydractinia

Homeobox genes are a large superfamily of protein-coding genes that encode for a 60 amino acid helix-turn-helix domain called the homeodomain (Holland 2013). Most homeobox genes are DNA-binding transcription factors (Holland 2013) that play key roles in early embryogenesis (Driever and Nüsslein-Volhard 1988), patterning (Pearson et al. 2005), development of the nervous system and sensory organs (Schulte and Frank 2014), and maintenance of embryonic stem cells (Young 2011). We identified 71 homeodomain-containing genes in the *H. symbiolongicarpus* genome and 82 in the *H. echinata* genome. Phylogenetic (Figures S24-25, Files S49-54) and secondary domain architecture-based approaches were able to resolve the ANTP, CERS, LIM, POU, PRD, SINE, and TALE homeobox classes, with a small number of genes remaining unclassified (Table S26). In both species, the ANTP-class homeodomains were the most abundant. Overall, *H. echinata* has 11 more homeobox genes than *H. symbiolongicarpus*, with expansions in the CERS, LIM, POU and PRD classes (Table S26). Four unclassified homeobox genes are unique to *H. echinata*. It is possible that some of these expansions in *H. echinata* may be duplicates from different alleles of the same gene that were not properly phased during the separation of haplotypes during the assembly process. All seven unclassified genes in *H. symbiolongicarpus* have a homolog to an unclassified gene in *H. echinata* (Table S27). Class-based annotation of homeodomain-containing genes based on phylogenetics, secondary domain information, and associated results from Orthofinder2 for both *Hydractinia* species can be found in Table S27.

### The HOX-L subclass of homeodomains in Hydractinia

Some of the most interesting genes to evolutionary biologists are those belonging to the Hox families of homeobox genes (Procino 2016). Hox genes are members of the ANTP class of homeoboxes, along with the Hox-like (‘extended Hox’) genes *Eve*, *Meox/Mox*, *Mnx,* and *Gbx*; the ParaHox cluster of *Gsx*, *Cdx*, and *Pdx/Xlox*; and the NK-like gene subclass (Holland 2013; Holland et al. 2007). The ANTP class is the largest and most diverse class, consisting of over 50 families; 37 of these families containing over 100 genes have been identified in humans (Holland et al. 2007). Hox and ParaHox genes are thought to have emerged prior to animal evolution and were subsequently lost, reduced, or absent in early-emerging taxa (Mendivil Ramos et al. 2012; Steinworth et al. 2023). In many bilaterians, Hox genes are arranged in at least one chromosomal cluster (Duboule 2007). Genomic linkage between Hox genes is present in extant cnidarians, although linked Hox and ParaHox genes were not found in previous cnidarian genome studies (Chapman et al. 2010; Putnam et al. 2007; Khalturin et al. 2019; Leclère et al. 2019; Gold et al. 2019; Jeon et al. 2019; DuBuc et al. 2012).

Both *Hydractinia* species possess several genes that belong to the HOX-L subclass (Figures S26-S27, Files S55-S60). These include several non-anterior (CenPost) cnidarian Hox genes, the ParaHox genes *Gsx* and *Cdx*, and the Hox-extended group *Mox*. Interestingly, the anterior Hox genes *Hox1* and *Hox2/Gsx-like* genes are absent in both species even though these genes have been found in other cnidarians, including hydrozoans (Chiori et al. 2009; Ryan et al. 2006). Additional members of the HOX-L repertoire that are present in other cnidarians but are absent in *Hydractinia* are genes encoding for the Hox-extended gene *Eve* and the ParaHox genes *Pdx/Xlox* (Leclère et al. 2019; Gold et al. 2019; Ryan et al. 2006). A primitive Hox cluster has been observed in anthozoan cnidarians but has not been found in hydrozoans (Chourrout et al. 2006; DuBuc et al. 2012). However, there appears to be some linkage of Hox genes in both *Hydractinia* genomes (Figure 3). This includes linkage of several cnidarian-specific Hox genes in *H. symbiolongicarpus* and linkage of a cnidarian Hox gene with the ParaHox gene *Gsx* in both *Hydractinia* species. Interestingly, the linkage of a Hox and ParaHox gene has not been shown in any other cnidarian genome. A comparison of phylogenetic relatedness and synteny analysis of various cnidarian species suggests that *Hydractinia* species likely lost the HOX-L genes *Hox1* and *Eve* (Figure 3). These genes are clustered together in anthozoans (DuBuc et al. 2012; Zimmermann et al. 2022) and *Eve* is found in close proximity to human Hox clusters (Faiella et al. 1991; D’Esposito et al. 1991). Interestingly, *Hydra* has retained a *Hox1* homolog but has also lost *Eve* (Putnam et al. 2007). Finally, the two *Hydractinia* genomes show for the first time that bilaterian-like ParaHox genes may have once been located near the central/posterior region of the Hox cluster (Figure 3). Therefore, the last common ancestor of cnidarians presumably had a linked Hox/ParaHox cluster (Figure 3) flanked by NK-class and other homeobox genes (D’Esposito et al. 1991). This could highlight that breaking apart the Hox and ParaHox cluster (which occurred in the bilaterian ancestor) was instrumental for their evolution.

**Figure 3:**
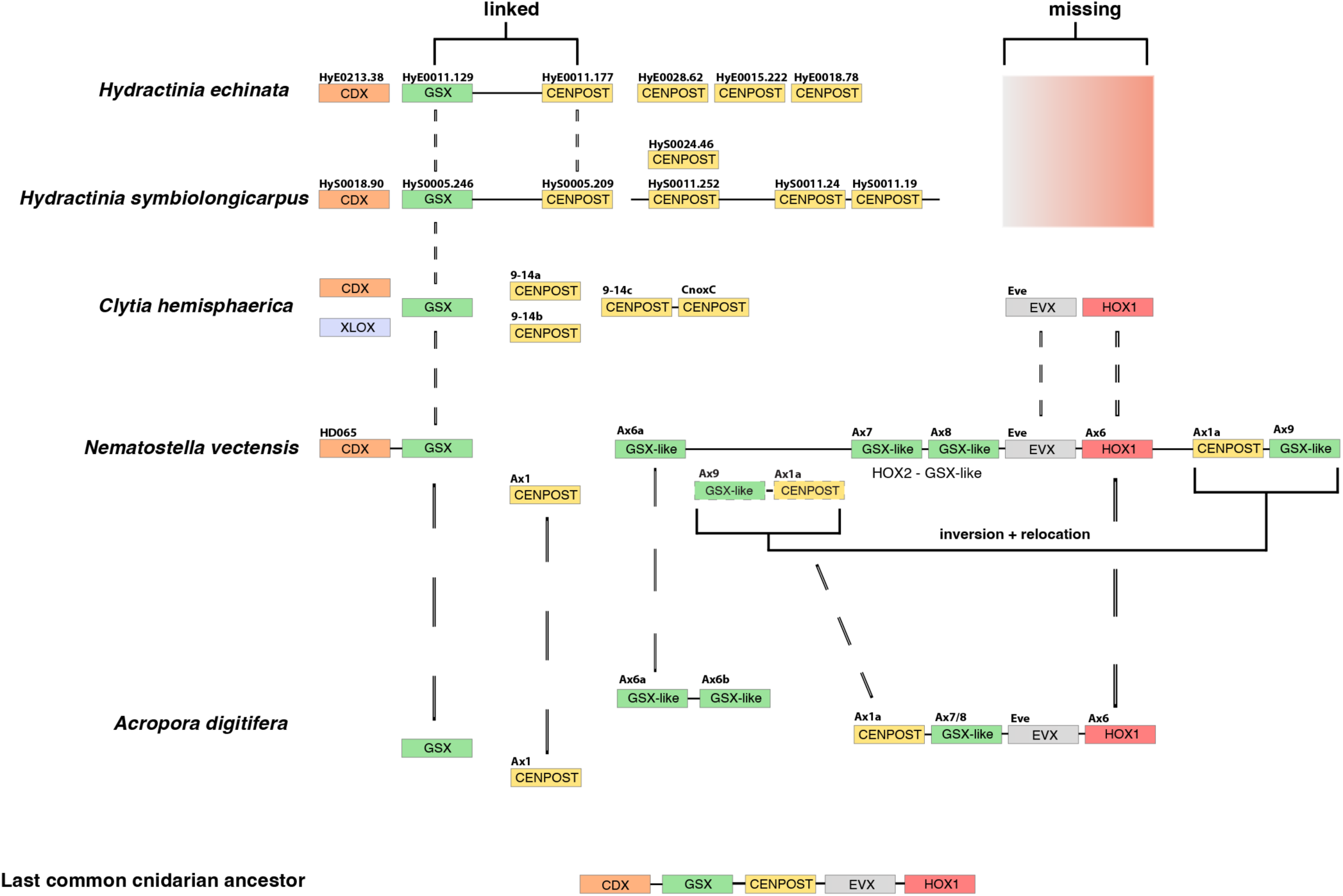
Genomic organization of Hox and ParaHox genes in five cnidarian genomes. Solid lines sharing homeobox genes represent genomic scaffolds. Scaffold and gene ID numbering in *Hydractinia* genomes is shown above gene boxes. Broken lines depict homologous cnidarian-specific Hox genes. Alternative gene names are shown above gene boxes for *C. hemisphaerica*, *N. vectensis*, and *A. digitifera*.

To determine the spatial patterning role of some of the homeobox genes relative to other known expression patterns, we performed colorimetric RNA *in situ* hybridization. Expression patterns for a subset of Hox genes at different stages of *Hydractinia*’s life cycle were determined (Figure S28). Overall, a number of genes show a somatic patterning role during early larva formation, while other Hox genes are maternally expressed during sexual development. This suggests that Hox genes may have an important role in egg formation.

### The Allorecognition Complex

Allorecognition is controlled by at least two linked genes, *Allorecognition 1* (*Alr1*) and *Allorecognition 2* (*Alr2*) in *Hydractinia* (Nicotra 2019; Nicotra et al. 2009; Rosa et al. 2010). Both encode single-pass transmembrane proteins with highly polymorphic extracellular domains, with the allorecognition response being controlled by whether colonies share alleles at these loci. In previous work, we examined the partially assembled genome of a strain of *H. symbiolongicarpus* that is homozygous at *Alr1* and *Alr2* and discovered that both genes are part of a family of immunoglobulin superfamily genes that reside in a genomic interval called the Allorecognition Complex (ARC) (Huene et al. 2022). We identified *Alr1* and *Alr2* on separate scaffolds within the *H. symbiolongicarpus* reference genome, as well as a second *Alr1* allele on a third scaffold. These alleles were likely retained in the final assembly because they were sufficiently divergent from each other not to be recognized as alleles of the same gene. We identified 19 additional genes predicted to encode full-length Alr proteins similar to those previously described (Huene et al. 2022), as well as 44 gene models with some sequence similarity to Alr1 or Alr2 that were not predicted to encode cell surface proteins, suggesting they were pseudogenes. Within the genome, most of these Alr1/Alr2-like gene models are located in four clusters (Figure S29). Additional work will be required to phase these contigs into two ARC haplotypes and assign orthology between them and the Alr genes already identified (Huene et al. 2022). Subsequent work focused on elucidating the genomic structure of the ARC based on these sequence data has shown that there are 41 *Alr-*-like loci in this region, with more than half of these genes located within one of three Alr clusters. While the individual Alr proteins encoded by these genes have low overall sequence identity, the domain architecture of these proteins, along with structure-based predictions using AlphaFold, confirm that these Alr proteins are members of the immunoglobulin superfamily (Huene et al. 2022).

### Single-cell transcriptomics of adult animals

A critical part of establishing *Hydractinia* as a useful research organism is having a list of cell-type specific markers for all cell types in the adult animal. Single cell transcriptome analysis of adult *H. symbiolongicarpus* 291-10 male animals was performed using the 10X Genomics platform (detailed methods in Supplement). Briefly, cell suspensions of dissociated adult feeding and sexual polyps and associated connective mat tissue were prepared and two samples were resuspended in different final buffers (3XPBS or calcium– and magnesium-free seawater minus EGTA) followed by subsequent 10X single-cell library construction. These two libraries were sequenced using the Illumina NovaSeq 6000 sequencing system. Statistics from each library can be found in Table S28. The two libraries were ultimately combined after analyzing them separately (Figure S30) and determining that they were very similar. Downstream analyses of these sequence data was performed with both the 10X Cell Ranger pipeline version 7.0.1 and the R package Seurat version 4.3.0 (Satija et al. 2015), ultimately yielding heatmaps and UMAP plots for the visualization of cell clusters (detailed methods in Supplement, File S61). The final clustering after filtering of technical artifacts (primarily removing sperm captured with another cell, termed ‘sperm doublets’; see Supplement and File S61) with Seurat resulted in 18 clusters from a total of 8,888 cells (Figure 4A). A heatmap was generated to show top variable ‘marker’ genes for each cluster (Figure S31).

**Figure 4:**
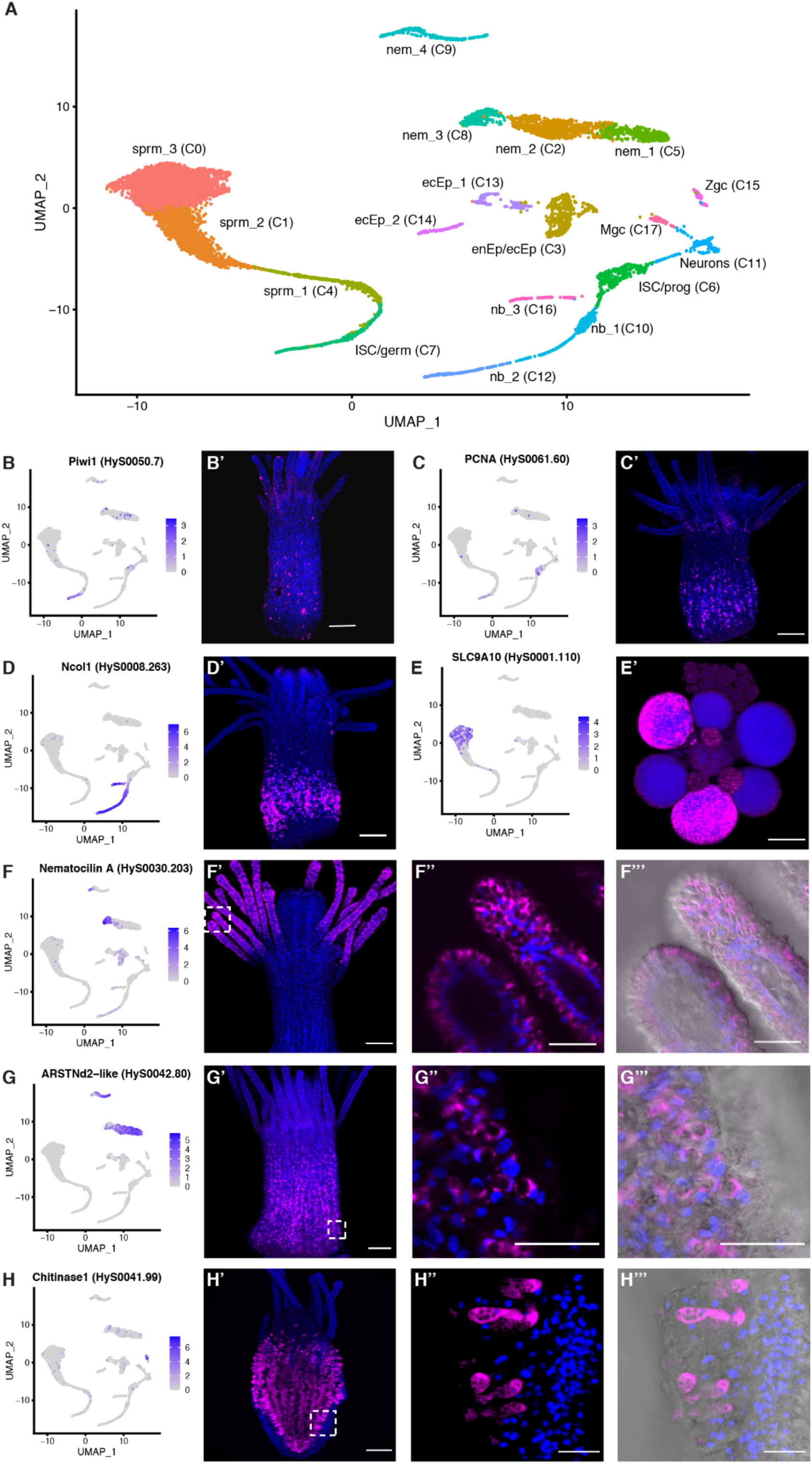
H*y*dractinia single-cell atlas represented as a labeled UMAP and validation of several cell type markers using fluorescent in situ hybridization (FISH). (A) *Hydractinia* single-cell atlas UMAP with 18 clusters (C0-C17). (B-F) UMAP expression of select marker genes (left panels) and spatial expression pattern of marker gene in polyps via FISH (right panel). Blue staining = Hoechst, Pink = marker gene. *Piwi1* (B) and *PCNA* (C) expression in the i-cell band in the middle of the body column of a feeding polyp. (D) *Ncol1* expression in nematoblasts in the lower body column of a feeding polyp. (E) *SLC9A10* expression in mature sperm cells in gonads of male sexual polyps. (F) *Nematocilin A* expression in a subset of nematocytes in the tentacles of a feeding polyp. Close up of tentacles in panels 3 and 4 both show higher magnification images from the same polyp as in panel 2, showing expression is specific to cnidocytes. Panel 4 adds DIC. (G) *ARSTNd2-like* expression in a subset of nematocytes in the body column of a feeding polyp. Panels 3 and 4 both show higher magnification images from the same polyp as in panel 2, showing expression is specific to cnidocytes. Panel 4 adds DIC. (H) *Chitinase 1* expression in gland cells in the endodermal epithelial cell layer of a feeding polyp. Panel 3 and 4 both show higher magnification images from the same polyp as in panel 2, showing expression is specific to gland cells. Panel 4 adds DIC. All images shown were projected from confocal stacks. All scale bars = 100 µm. Abbreviations in (A): ecEP: ectodermal epithelial cell, enEP: endodermal epithelial cell, germ: germ cell, ISC: interstitial stem cell, Mgc: mucous gland cell, nb: nematoblast, nem: nematocyte, prog: progenitor, sprm: sperm, Zgc: zymogen gland cell.

Each cluster was then classified as a putative cell type or cell state through the annotation of these marker genes; these included distinct clusters of ectodermal (epidermal) and endodermal (gastrodermal) epithelial cells, mucous and zymogen gland cells, neurons, nematoblasts, nematocytes, germ cells, developing stages of sperm, and two clusters of i-cells (Figure 4A). These i-cell clusters probably include early progenitor cells as puripotent i-cells are a rare population (Varley et al. 2023; Chrysostomou et al. 2022; DuBuc et al. 2020), thus we have labeled them as ISC/prog on our UMAP. UMAP expression patterns for individual genes that were used to identify and annotate the clusters based on previous literature can be found in Figure S32 and further details are provided in Table S29. We grouped these clusters into seven major cell ‘types’: Sperm and Spermatocytes (clusters C0, C1, and C4), Nematocytes (C2, C5, C8, and C9), Epithelial Cells (C3, C13, and C14), i-cells/Germ Cells (C6 and C7), Nematoblasts (C10, C12, and C16), Neurons (C11), and Gland Cells (C15 and C17).

A subset of seven different cell-type marker genes were chosen for fluorescence *in situ* hybridization (FISH) for validation and to visualize spatial expression patterns of various cell types in adult polyps (Figure 4B-H), including two genes that have been previously published for *Hydractinia* [*Piwi1* for marking i-cells/progenitors and *Ncol1* for marking all stages of maturing nematoblasts (Bradshaw et al. 2015)]. The five remaining genes can be considered new cell-type markers for *Hydractinia*. We observed that the proliferating cell nuclear antigen *PCNA*, a known proliferation and broad stem cell marker in other animals (Wagner et al. 2011), marks cells present in the i-cell band; *SLC9A10*, a member of the sodium-hydrogen exchanger (NHE) family required for male fertility and sperm motility (Wang et al. 2003) marks mature sperm in gonads of male sexual polyps; *nematocilin A*, a known structural component of the cnidocil mechanosensory cilium trigger of mature cnidocytes in *Hydra* (Hwang et al. 2008), marks mature cnidocytes in tentacles; *ARSTNd2-like* (previously undescribed) marks cnidocytes in the polyp body column, and *Chitinase1*, a gland/secretory cell marker in cnidarians (Klug et al. 1984; Sebé-Pedrós et al. 2018) marks endodermal gland cells. These results represent a significant step towards defining the major cell types in *Hydractinia* and the gene expression patterns that define them. A list of all cluster marker genes according to cell type from the Seurat analysis can be found in Table S30.

We then explored the evolutionary profile of marker genes from the 18 individual clusters and the seven cell types (split further into nine groups, Figure 5A) using strict filtering criteria (detailed methods in Supplement). We found that, compared with other cell types (and clusters), i-cells and progenitors (ISC/prog cluster 6, 5.3% lineage-specific; ISC/germ cluster 7, 12.5% lineage-specific; all i-cells and progenitors, 9.5% lineage-specific) and early spermatogonia (cluster 4, 9.7% lineage-specific) are defined primarily by genes that are shared with other animals rather than lineage-specific genes, providing evidence that the toolkit employed by these cell types has a shared ancestry with other animals (Figure 5A). Nematoblasts and nematocytes – cell types that are specific to cnidarians – were marked by a high proportion of phylum-specific or within-phylum genes (nematoblasts 49%, nematocytes 32.5%), which was expected. Further probing into the i-cell cluster profile (clusters C6 and 7) to analyze how widespread the i-cell/progenitor marker genes were among animals in our dataset, we plotted how many species in our orthology-inference analysis shared each i-cell marker gene and found that the vast majority of the genes that mark i-cells are present in 40 or more species (Figure 5B). Overall, our finding that the 317 i-cell marker genes were widely shared among all animals was surprising, given that the *H. symbiolongicarpus* genome has a higher proportion of phylum-specific and within-Cnidaria-specific genes (23%) than any of the other 41 animals in our orthology-inference analysis. The *Hydractinia* single cell dataset has an even higher proportion of phylum-specific and within-Cnidaria-specific genes (30.8%). This finding indicates that the stem cells of this cnidarian may use a widely shared gene toolkit found in nearly all animals that holds promise for future exploration. Further study of these i-cell marker genes will result in knowledge that will likely help aid in understanding evolutionarily conserved functions of stem cells in other animals. It remains to be seen whether other animals share the same or partially overlapping toolkit of genes in their own stem cells, an important question that is beginning to be addressed using new methodologies currently under development (Wang et al. 2021).

**Figure 5:**
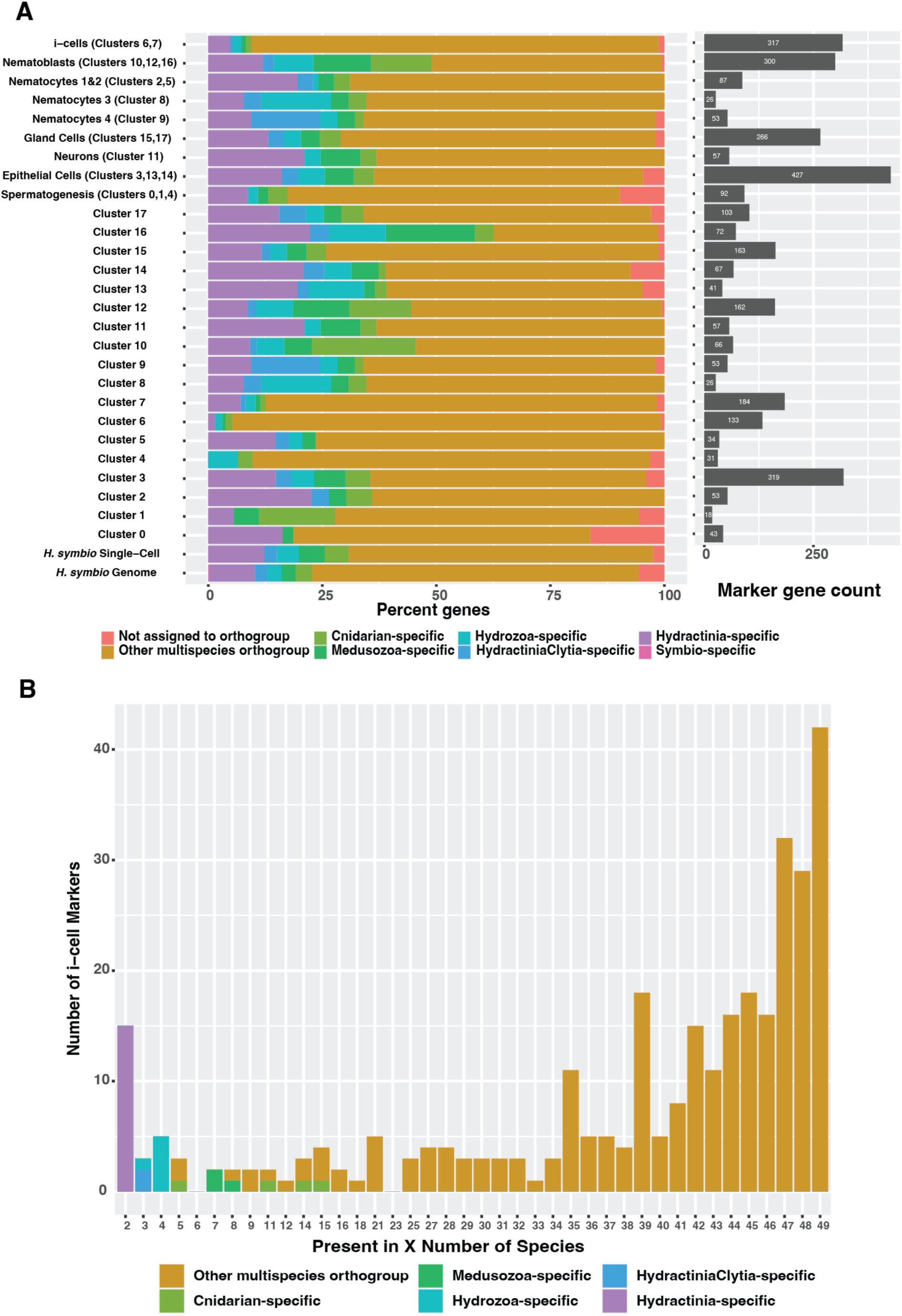
Results from the lineage-specificity analysis using OrthoFinder2 results and the UMAP cluster marker genes. (A) Stacked bar chart showing the percentage of *H. symbiolongicarpus* single-cell atlas cluster markers shared among animal phyla. The bottom legend shows eight different categories, dividing the markers into different groups depending on how the orthologs are shared among the species. “Not assigned to orthogroup’’ represents markers that could not be placed into an orthogroup. The other categories are markers that have at least one homolog between *H. symbiolongicarpus* and that category, except for the “Symbio-specific” category which represents markers that fell in orthogroups with only *H. symbiolongicarpus* genes. For example, hypothetical marker gene A from *H. symbiolongicarpus* would be an “Other multispecies orthogroup” marker if it was found in *H. symbiolongicarpus* and at least one animal outside of cnidaria, but it would be a “Cnidarian-specific” marker if it was found in *H. symbiolongicarpus* and at least one cnidarian outside medusozoa. Stacked bars represent the seven major cell types split into nine groups, followed by all individual clusters, and finally the total genes expressed in the *Hydractinia* single-cell dataset (16,069 genes) and total genes predicted from the *Hydractinia* genome (22,022 genes). The marker gene count bars on the right indicate how many markers are present in each major cell type and cluster. (B) Histogram dividing the 317 orthogroup-assigned i-cell (clusters 6 and 7) markers by how many are shared by X number of species. Legend is the same as in (A) but the following categories are excluded from this chart: unassigned genes (two genes) and *H. symbiolongicarpus*-specific genes (none).

## DISCUSSION

Understanding the functions and evolutionary context of genes that underlie complex processes like regeneration and cell-type specification requires a broad approach. In this study, we used two newly sequenced *Hydractinia* genomes to perform an orthology analysis with a broad sampling of 43 animal genomes and six related outgroup species to better understand the evolutionary relationships of their genes, especially the genes that define *Hydractinia*’s i-cells and progenitors. By viewing these genes through the lens of orthology across diverse animal phyla, and by leveraging single-cell RNAseq data, we revealed that *Hydractinia* stem cells and progenitors use a widely-conserved toolkit of genes. This makes *Hydractinia* a promising model organism for future exploration of stem cell biology and regenerative medicine, and enables us to link discoveries in *Hydractinia* to other animals. As research into more diverse model species is undertaken, this approach can enable us to gain a deeper understanding of how widely-spread genes that underlie complex biological processes truly are.

## MATERIALS and METHODS

### Genome sequencing and assembly

Genomic DNA was prepared from adult polyps from a single strain for each species (291-10 males for *H. symbiolongicarpus* and F4 females from *H. echinata*). PacBio long-read and Illumina short-read data were generated. Canu was used as the contig assembler. Scaffolding was done by Dovetail HiRise scaffolding with Illumina Chicago libraries.

### Gene model prediction and annotation

Gene models were generated with a pipeline that involved both PASA and Augustus. Strand-specific RNAseq data from each species was used as input at different points of the pipeline as reads and as assembled transcripts. Functional annotation was performed with a DIAMOND search of NCBI’s nr database and PANNZER2.

### Orthology inference, phylogenetic analyses, and divergence time estimates

Orthology-inference analysis was performed on a splice-filtered proteome dataset of 49 species from 15 metazoan phyla and four non-metazoan outgroups. Orthology assignment was performed using OrthoFinder version 2.2.7. Divergence times between *H. echinata* and *H. symbiolongicarpus* and between other cnidarian lineages were estimated by inferring a time-calibrated maximum-likelihood phylogeny using only single copy orthologs. The topology of our maximum likelihood phylogenetic tree was inferred using IQ-Tree2, and divergence date estimates were calculated for major nodes on the tree using a Langley-Fitch approach together with the TN algorithm, using r8s version 1.8.1 (Figure 1B).

### Orthogroup lineage specificity and overall patterns

Output from Orthofinder was processed using custom R scripts (Files S25-S27) to analyze patterns of presence and absence of orthogroups across taxa and characterize the taxon-specificity of each orthogroup. Taxon specificity and other related information for each *H. symbiolongicarpus* and *H. echinata* gene model can be found in Supplemental Table S11 tabs ‘X.10’ and ‘X.11’.

### Estimating the evolutionary dynamics of gene families using CAFÉ

We used the software package CAFE v. 4.2.1 to estimate ancestral gene family sizes and changes in gene family size among 15 cnidarian species, as well as to infer which gene families are significantly faster evolving in specific cnidarian lineages. As input, we provided our time-calibrated tree and the gene counts per species for a subset of the orthogroups inferred by Orthofinder.

### Single-cell transcriptomics of adult animals and OrthoMarker analyses

Tissue from adult male *H. symbiolongicarpus* clone 291-10 was dissociated in 1% Pronase E in calcium– and magnesium-free artificial seawater (CMFASW) with EGTA for 90 minutes total. The cell suspension was filtered through a 70µm Flowmi cell filter, pelleted at 300 rcf for 5 minutes at 4C, and the pellet gently resuspended in either CMFASW without EGTA or 3XPBS. This cell suspension was filtered through a 40µm Flowmi cell filter and placed on ice. 10X single cell 3’ version 3 RNAseq library construction was performed at the University of Florida’s Interdisciplinary Center for Biotechnology Research. Libraries were sequenced at the NIH Intramural Sequencing Center using the Illumina NovaSeq 6000_SP sequencing system. The 10X Cell Ranger pipeline version 7.0.1 was used to pre-process the sequencing data for downstream analysis. The R package Seurat version 4.3.0 was used to generate clusters, find marker genes for each cluster, and further analyze the data. A marker gene list for each cluster was created using Seurat and the settings used in Siebert et al. 2019. The Orthofinder results (Files S8-S14) were used to apply several levels of taxon-specificity to the marker gene list using R and the ‘dyplr’ package. The R package ‘ggplot’ was used to create the bar plot and histogram shown as Figure 5a and Figure 5b, respectively. Markers were validated with fluorescence in situ hybridization (see Supplement) and primers for those genes are found in Table S31.

## Data access

This Whole Genome Shotgun project has been deposited at DDBJ/ENA/GenBank under the accession JARYZW000000000 (*H. symbiolongicarpus*) and JASGCC000000000 (*H. echinata*). The version described in this paper is version JARYZW010000000 (*H. symbiolongicarpus*) and JASGCC010000000 (*H. echinata*). All sequencing read data related to this project can be found in the SRA under BioProject PRJNA807936 (*H. symbiolongicarpus*) and PRJNA812777 (*H. echinata*). The *Hydractinia* genome project portal: https://research.nhgri.nih.gov/hydractinia/ provides a rich source of data for both species, including a BLAST interface, genome browser, DNA and protein sequence downloads, and functional annotation of gene models, as well as a single cell browser and RNA-Seq expression data.

## Acknowledgments

We would like to thank the following people: J. Spencer Johnston of Texas A&M University for propidium iodine-based genome size estimation. Rob Steele for providing *Hydra vulgaris* strain 105. Alice Young and many others at the NIH Intramural Sequencing Center (NISC) for DNA and RNA sequencing library construction, sequencing, as well as advice and support. Dovetail Genomics for providing Chicago libraries and scaffolding services. Leo Buss for advice and support. Gunter Plickert and Philipp Schiffer for early RNA-seq datasets not included in the final paper. Bernie Koch, Steve Bond, and Derek Gildea for advice and help. Suiyuan Zhang for making data available on the *Hydractinia* genome portal. Alexandrea Duscher for creating the *Chit1* riboprobe and assisting with the single-cell experiment. Mackenzie Simon-Collins for helping to optimize the cell dissociation protocol. Malcolm Maden for providing bench space during our single-cell experiment. Yanping Zhang and Alex Deiulio at the UF ICBR Gene Expression & Genotyping Core (RRID:SCR_019145) for advice and for constructing the 10X single-cell libraries. This work utilized the Biowulf high-performance supercomputing resource of the Center for Information Technology at the National Institutes of Health (https://hpc.nih.gov).

## Author contributions

C.E.S., P.C., M.N., U.F, and A.D.B. conceived the study; C.E.S., E.S.C., and A.D.B. wrote the paper with revisions by G.Q-A., T.Q.D., D.D.J., E.P.N., T.G.W., O.S., M.N., and U.F.; J.M.G, S.M.S., and F. provided samples for sequencing; C.E.S., S.N.B., P.G., S.Koren, and A.P. performed whole genome and bulk and single-cell RNA-seq sequencing and assembly; C.E.S., A.-D.N., S.N.B., and P.G. built gene models and provided genome annotation; E.S.C., W.Y.W., and P.G. performed phylogenetic and orthofinder analyses; C.E.S., J.W., G.Q.A., J.G., D.D.J, B.B., and A.W. performed head regeneration and single-cell RNAseq experiments and analysis; D.D.J., J.W., G.Q-A., and T.Q.D. performed ISH experiments and imaging; C.E.S, E.S.C, J.W., G.Q.A., W.Y.W., A-D.N., S.N.B., L.D., P.G., S.Koren, T.Q.D., E.P.N., S.Klasfeld, S.G.G., A.P., J.M., and O.S. processed raw data and conducted data analysis; A.-D.N., R.T.M., and T.G.W. contributed new analytic tools/resources and additional data.

## REFERENCES

1. Beagley CT, Okimoto R, Wolstenholme DR. 1998. The mitochondrial genome of the sea anemone Metridium senile (Cnidaria): introns, a paucity of tRNA genes, and a near-standard genetic code. Genetics 148: 1091–108.

2. Benson G. 1999. Tandem repeats finder: a program to analyze DNA sequences. Nucleic Acids Research 27: 573–580.

3. Bermudez-Santana C, Attolini CS-O, Kirsten T, Engelhardt J, Prohaska SJ, Steigele S, Stadler PF. 2010. Genomic organization of eukaryotic tRNAs. BMC Genomics 11: 270.

4. Bradshaw B, Thompson K, Frank U. 2015. Distinct mechanisms underlie oral vs aboral regeneration in the cnidarian Hydractinia echinata ed. A. Sánchez Alvarado. eLife 4: e05506.

5. Bridge D, Cunningham CW, Schierwater B, DeSalle R, Buss LW. 1992. Class-level relationships in the phylum Cnidaria: evidence from mitochondrial genome structure. Proc Natl Acad Sci U S A 89: 8750–3.

6. Brugler MR, France SC. 2007. The complete mitochondrial genome of the black coral Chrysopathes formosa (Cnidaria:Anthozoa:Antipatharia) supports classification of antipatharians within the subclass Hexacorallia. Mol Phylogenet Evol 42: 776–88.

7. Buchfink B, Xie C, Huson DH. 2015. Fast and sensitive protein alignment using DIAMOND. Nature Methods 12: 59–60.

8. Chan PP, Lin BY, Mak AJ, Lowe TM. 2021. tRNAscan-SE 2.0: improved detection and functional classification of transfer RNA genes. Nucleic Acids Research 49: 9077–9096.

9. Chapman JA, Kirkness EF, Simakov O, Hampson SE, Mitros T, Weinmaier T, Rattei T, Balasubramanian PG, Borman J, Busam D, et al. 2010. The dynamic genome of Hydra. Nature 464: 592–596.

10. Chen C, Chiou CY, Dai CF, Chen CA. 2008. Unique mitogenomic features in the scleractinian family pocilloporidae (scleractinia: astrocoeniina). Mar Biotechnol (NY) 10: 538–53.

11. Chen R, Sanders SM, Ma Z, Paschall J, Chang ES, Riscoe BM, Schnitzler CE, Baxevanis AD, Nicotra ML. 2023. XY sex determination in a cnidarian. BMC Biology 21: 32.

12. Chin C-S, Alexander DH, Marks P, Klammer AA, Drake J, Heiner C, Clum A, Copeland A, Huddleston J, Eichler EE, et al. 2013. Nonhybrid, finished microbial genome assemblies from long-read SMRT sequencing data. Nature Methods 10: 563–569.

13. Chin C-S, Peluso P, Sedlazeck FJ, Nattestad M, Concepcion GT, Clum A, Dunn C, O’Malley R, Figueroa-Balderas R, Morales-Cruz A, et al. 2016. Phased diploid genome assembly with single-molecule real-time sequencing. Nat Methods 13: 1050–1054.

14. Chiori R, Jager M, Denker E, Wincker P, Silva CD, Guyader HL, Manuel M, Quéinnec E. 2009. Are Hox Genes Ancestrally Involved in Axial Patterning? Evidence from the Hydrozoan Clytia hemisphaerica (Cnidaria). PLOS ONE 4: e4231.

15. Chourrout D, Delsuc F, Chourrout P, Edvardsen RB, Rentzsch F, Renfer E, Jensen MF, Zhu B, de Jong P, Steele RE, et al. 2006. Minimal ProtoHox cluster inferred from bilaterian and cnidarian Hox complements. Nature 442: 684–687.

16. Chrysostomou E, Flici H, Gornik SG, Salinas-Saavedra M, Gahan JM, McMahon ET, Thompson K, Hanley S, Kincoyne M, Schnitzler CE, et al. 2022. A cellular and molecular analysis of SoxB-driven neurogenesis in a cnidarian ed. M.E. Bronner. eLife 11: e78793.

17. Cloix C, Tutois S, Mathieu O, Cuvillier C, Espagnol MC, Picard G, Tourmente S. 2000. Analysis of 5S rDNA Arrays in Arabidopsis thaliana: Physical Mapping and Chromosome-Specific Polymorphisms. Genome Res 10: 679–690.

18. Darrow EM, Chadwick BP. 2014. A novel tRNA variable number tandem repeat at human chromosome 1q23.3 is implicated as a boundary element based on conservation of a CTCF motif in mouse. Nucleic Acids Research 42: 6421–6435.

19. De Bie T, Cristianini N, Demuth JP, Hahn MW. 2006. CAFE: a computational tool for the study of gene family evolution. Bioinformatics 22: 1269–1271.

20. Derelle R, Momose T, Manuel M, Silva CD, Wincker P, Houliston E. 2010. Convergent origins and rapid evolution of spliced leader trans-splicing in Metazoa: Insights from the Ctenophora and Hydrozoa. RNA 16: 696–707.

21. D’Esposito M, Morelli F, Acampora D, Migliaccio E, Simeone A, Boncinelli E. 1991. EVX2, a human homeobox gene homologous to the even-skipped segmentation gene, is localized at the 5ʹ end of HOX4 locus on chromosome 2. Genomics 10: 43–50.

22. Driever W, Nüsslein-Volhard C. 1988. The bicoid protein determines position in the Drosophila embryo in a concentration-dependent manner. Cell 54: 95–104.

23. Duboule D. 2007. The rise and fall of Hox gene clusters. Development 134: 2549–2560.

24. DuBuc TQ, Ryan JF, Shinzato C, Satoh N, Martindale MQ. 2012. Coral Comparative Genomics Reveal Expanded Hox Cluster in the Cnidarian–Bilaterian Ancestor. Integrative and Comparative Biology 52: 835–841.

25. DuBuc TQ, Schnitzler CE, Chrysostomou E, McMahon ET, Febrimarsa, Gahan JM, Buggie T, Gornik SG, Hanley S, Barreira SN, et al. 2020. Transcription factor AP2 controls cnidarian germ cell induction. Science 367: 757–762.

26. English AC, Richards S, Han Y, Wang M, Vee V, Qu J, Qin X, Muzny DM, Reid JG, Worley KC, et al. 2012. Mind the Gap: Upgrading Genomes with Pacific Biosciences RS Long-Read Sequencing Technology. PLOS ONE 7: e47768.

27. Faiella A, D’Esposito M, Rambaldi M, Acampora D, Balsfiore S, Stornaiuolo A, Mallamaci A, Migliaccio E, Gulisano M, Simeone A, et al. 1991. Isolation and mapping of EVx1, a human homeobox gene homologus to even-skipped, localized at the 5ʹ end of Hox1 locus on chromosome 7. Nucleic Acids Research 19: 6541–6545.

28. Friedländer MR, Mackowiak SD, Li N, Chen W, Rajewsky N. 2012. miRDeep2 accurately identifies known and hundreds of novel microRNA genes in seven animal clades. Nucleic Acids Research 40: 37–52.

29. Gold DA, Katsuki T, Li Y, Yan X, Regulski M, Ibberson D, Holstein T, Steele RE, Jacobs DK, Greenspan RJ. 2019. The genome of the jellyfish Aurelia and the evolution of animal complexity. Nature Ecology & Evolution 3: 96–104.

30. Guo L, Accorsi A, He S, Guerrero-Hernández C, Sivagnanam S, McKinney S, Gibson M, Sánchez Alvarado A. 2018. An adaptable chromosome preparation methodology for use in invertebrate research organisms. BMC Biol 16: 25.

31. Haas BJ, Salzberg SL, Zhu W, Pertea M, Allen JE, Orvis J, White O, Buell CR, Wortman JR. 2008. Automated eukaryotic gene structure annotation using EVidenceModeler and the Program to Assemble Spliced Alignments. Genome Biol 9: R7.

32. Han MV, Thomas GWC, Lugo-Martinez J, Hahn MW. 2013. Estimating Gene Gain and Loss Rates in the Presence of Error in Genome Assembly and Annotation Using CAFE 3. Molecular Biology and Evolution 30: 1987–1997.

33. Hare EE, Johnston JS. 2011. Genome Size Determination Using Flow Cytometry of Propidium Iodide-Stained Nuclei. In Molecular Methods for Evolutionary Genetics (eds. V. Orgogozo and M.V. Rockman), *Methods in Molecular Biology*, pp. 3–12, Humana Press, Totowa, NJ 10.1007/978-1-61779-228-1_1 (Accessed May 4, 2021).

34. Hastings KEM. 2005. SL trans-splicing: easy come or easy go? Trends in Genetics 21: 240–247.

35. Holland PW, Booth HAF, Bruford EA. 2007. Classification and nomenclature of all human homeobox genes. BMC Biology 5: 47.

36. Holland PWH. 2013. Evolution of homeobox genes. WIREs Developmental Biology 2: 31–45.

37. Huene AL, Sanders SM, Ma Z, Nguyen A-D, Koren S, Michaca MH, Mullikin JC, Phillippy AM, Schnitzler CE, Baxevanis AD, et al. 2022. A family of unusual immunoglobulin superfamily genes in an invertebrate histocompatibility complex. Proceedings of the National Academy of Sciences 119: e2207374119.

38. Hwang JS, Takaku Y, Chapman J, Ikeo K, David CN, Gojobori T. 2008. Cilium Evolution: Identification of a Novel Protein, Nematocilin, in the Mechanosensory Cilium of Hydra Nematocytes. Molecular Biology and Evolution 25: 2009– 2017.

39. Jeon Y, Park SG, Lee N, Weber JA, Kim H-S, Hwang S-J, Woo S, Kim H-M, Bhak Y, Jeon S, et al. 2019. The Draft Genome of an Octocoral, Dendronephthya gigantea. Genome Biology and Evolution 11: 949–953.

40. Kalvari I, Argasinska J, Quinones-Olvera N, Nawrocki EP, Rivas E, Eddy SR, Bateman A, Finn RD, Petrov AI. 2018. Rfam 13.0: shifting to a genome-centric resource for non-coding RNA families. Nucleic Acids Research 46: D335–D342.

41. Kayal E, Bentlage B, Cartwright P, Yanagihara AA, Lindsay DJ, Hopcroft RR, Collins AG. 2015a. Phylogenetic analysis of higher-level relationships within Hydroidolina (Cnidaria: Hydrozoa) using mitochondrial genome data and insight into their mitochondrial transcription. PeerJ 3: e1403.

42. Kayal E, Bentlage B, Cartwright P, Yanagihara AA, Lindsay DJ, Hopcroft RR, Collins AG. 2015b. Phylogenetic analysis of higher-level relationships within Hydroidolina (Cnidaria: Hydrozoa) using mitochondrial genome data and insight into their mitochondrial transcription. PeerJ 3: e1403.

43. Kayal E, Bentlage B, Collins AG, Kayal M, Pirro S, Lavrov DV. 2012. Evolution of linear mitochondrial genomes in medusozoan cnidarians. Genome Biol Evol 4: 1–12.

44. Kayal E, Lavrov DV. 2008. The mitochondrial genome of Hydra oligactis (Cnidaria, Hydrozoa) sheds new light on animal mtDNA evolution and cnidarian phylogeny. Gene 410: 177–86.

45. Khalturin K, Shinzato C, Khalturina M, Hamada M, Fujie M, Koyanagi R, Kanda M, Goto H, Anton-Erxleben F, Toyokawa M, et al. 2019. Medusozoan genomes inform the evolution of the jellyfish body plan. Nature Ecology & Evolution 3: 811–822.

46. Klug M, Tardent P, Smid I, Holstein T. 1984. Presence and localization of chitinase in Hydra and Podocoryne (Cnidaria, Hydrozoa). Journal of Experimental Zoology 229: 69–72.

47. Koren S, Walenz BP, Berlin K, Miller JR, Bergman NH, Phillippy AM. 2017. Canu: scalable and accurate long-read assembly via adaptive k-mer weighting and repeat separation. Genome Res 27: 722–736.

48. Kurtz S, Phillippy A, Delcher AL, Smoot M, Shumway M, Antonescu C, Salzberg SL. 2004. Versatile and open software for comparing large genomes. Genome Biol 5: R12.

49. Leclère L, Horin C, Chevalier S, Lapébie P, Dru P, Peron S, Jager M, Condamine T, Pottin K, Romano S, et al. 2019. The genome of the jellyfish Clytia hemisphaerica and the evolution of the cnidarian life-cycle. Nature Ecology & Evolution 3: 801–810.

50. Long EO, Dawid IB. 1980. Repeated Genes in Eukaryotes. Annual Review of Biochemistry 49: 727–764.

51. Maxwell EK, Schnitzler CE, Havlak P, Putnam NH, Nguyen A-D, Moreland RT, Baxevanis AD. 2014. Evolutionary profiling reveals the heterogeneous origins of classes of human disease genes: implications for modeling disease genetics in animals. BMC Evol Biol 14: 212.

52. Mendivil Ramos O, Barker D, Ferrier DEK. 2012. Ghost Loci Imply Hox and ParaHox Existence in the Last Common Ancestor of Animals. Current Biology 22: 1951–1956.

53. Moran Y, Praher D, Fredman D, Technau U. 2013. The Evolution of MicroRNA Pathway Protein Components in Cnidaria. Molecular Biology and Evolution 30: 2541–2552.

54. Munro C, Cadis H, Pagnotta S, Houliston E, Huynh J-R. 2023. Conserved meiotic mechanisms in the cnidarian Clytia hemisphaerica revealed by Spo11 knockout. Science Advances 9: eadd2873.

55. Nawrocki EP, Eddy SR. 2013. Infernal 1.1: 100-fold faster RNA homology searches. Bioinformatics 29: 2933–2935.

56. Nicotra ML. 2019. Invertebrate allorecognition. Current Biology 29: R463–R467.

57. Nicotra ML, Powell AE, Rosengarten RD, Moreno M, Grimwood J, Lakkis FG, Dellaporta SL, Buss LW. 2009. A Hypervariable Invertebrate Allodeterminant. Current Biology 19: 583–589.

58. Nong W, Cao J, Li Y, Qu Z, Sun J, Swale T, Yip HY, Qian PY, Qiu JW, Kwan HS, et al. 2020. Jellyfish genomes reveal distinct homeobox gene clusters and conservation of small RNA processing. Nat Commun 11: 3051.

59. Pearson JC, Lemons D, McGinnis W. 2005. Modulating Hox gene functions during animal body patterning. Nat Rev Genet 6: 893–904.

60. Praher D, Zimmermann B, Dnyansagar R, Miller DJ, Moya A, Modepalli V, Fridrich A, Sher D, Friis-Møller L, Sundberg P, et al. 2021. Conservation and turnover of miRNAs and their highly complementary targets in early branching animals. Proceedings of the Royal Society B: Biological Sciences 288: 20203169.

61. Procino A. 2016. Class I Homeobox Genes, “The Rosetta Stone of the Cell Biology”, in the Regulation of Cardiovascular Development. Current Medicinal Chemistry 23: 265–275.

62. Putnam NH, O’Connell BL, Stites JC, Rice BJ, Blanchette M, Calef R, Troll CJ, Fields A, Hartley PD, Sugnet CW, et al. 2016. Chromosome-scale shotgun assembly using an in vitro method for long-range linkage. Genome Res 26: 342–350.

63. Putnam NH, Srivastava M, Hellsten U, Dirks B, Chapman J, Salamov A, Terry A, Shapiro H, Lindquist E, Kapitonov VV, et al. 2007. Sea Anemone Genome Reveals Ancestral Eumetazoan Gene Repertoire and Genomic Organization. Science 317: 86–94.

64. Rosa SFP, Powell AE, Rosengarten RD, Nicotra ML, Moreno MA, Grimwood J, Lakkis FG, Dellaporta SL, Buss LW. 2010. Hydractinia Allodeterminant alr1 Resides in an Immunoglobulin Superfamily-like Gene Complex. Current Biology 20: 1122–1127.

65. Ryan JF, Burton PM, Mazza ME, Kwong GK, Mullikin JC, Finnerty JR. 2006. The cnidarian-bilaterian ancestor possessed at least 56 homeoboxes: evidence from the starlet sea anemone, Nematostella vectensis. Genome Biol 7: R64.

66. Sanderson MJ. 2003. r8s: inferring absolute rates of molecular evolution and divergence times in the absence of a molecular clock. Bioinformatics 19: 301–302.

67. Satija R, Farrell JA, Gennert D, Schier AF, Regev A. 2015. Spatial reconstruction of single-cell gene expression data. Nature Biotechnology 33: 495–502.

68. Schulte D, Frank D. 2014. TALE transcription factors during early development of the vertebrate brain and eye. Developmental Dynamics 243: 99–116.

69. Sebé-Pedrós A, Saudemont B, Chomsky E, Plessier F, Mailhé M-P, Renno J, Loe-Mie Y, Lifshitz A, Mukamel Z, Schmutz S, et al. 2018. Cnidarian Cell Type Diversity and Regulation Revealed by Whole-Organism Single-Cell RNA-Seq. Cell 173: 1520–1534.e20.

70. Shao Z, Graf S, Chaga OY, Lavrov DV. 2006. Mitochondrial genome of the moon jelly Aurelia aurita (Cnidaria, Scyphozoa): A linear DNA molecule encoding a putative DNA-dependent DNA polymerase. Gene 381: 92–101.

71. Simakov O, Bredeson J, Berkoff K, Marletaz F, Mitros T, Schultz DT, O’Connell BL, Dear P, Martinez DE, Steele RE, et al. 2022. Deeply conserved synteny and the evolution of metazoan chromosomes. Science Advances 8: eabi5884.

72. Simão FA, Waterhouse RM, Ioannidis P, Kriventseva EV, Zdobnov EM. 2015. BUSCO: assessing genome assembly and annotation completeness with single-copy orthologs. Bioinformatics 31: 3210–3212.

73. Smith DR, Kayal E, Yanagihara AA, Collins AG, Pirro S, Keeling PJ. 2012. First complete mitochondrial genome sequence from a box jellyfish reveals a highly fragmented linear architecture and insights into telomere evolution. Genome Biol Evol 4: 52–8.

74. Song S, Jiang F, Yuan J, Guo W, Miao Y. 2013. Exceptionally high cumulative percentage of NUMTs originating from linear mitochondrial DNA molecules in the Hydra magnipapillata genome. BMC Genomics 14: 447.

75. Stampar SN, Broe MB, Macrander J, Reitzel AM, Brugler MR, Daly M. 2019. Linear Mitochondrial Genome in Anthozoa (Cnidaria): A Case Study in Ceriantharia. Sci Rep 9: 6094.

76. Steele RE, David CN, Technau U. 2011. A genomic view of 500 million years of cnidarian evolution. Trends in Genetics 27: 7–13.

77. Steinworth BM, Martindale MQ, Ryan JF. 2023. Gene Loss may have Shaped the Cnidarian and Bilaterian Hox and ParaHox Complement ed. S.F. Valverde. Genome Biology and Evolution 15: evac172.

78. Stover NA, Steele RE. 2001. Trans-spliced leader addition to mRNAs in a cnidarian. PNAS 98: 5693–5698.

79. Tawari B, Ali IKM, Scott C, Quail MA, Berriman M, Hall N, Clark CG. 2008. Patterns of Evolution in the Unique tRNA Gene Arrays of the Genus Entamoeba. Molecular Biology and Evolution 25: 187–198.

80. Török A, Schiffer PH, Schnitzler CE, Ford K, Mullikin JC, Baxevanis AD, Bacic A, Frank U, Gornik SG. 2016. The cnidarian Hydractinia echinata employs canonical and highly adapted histones to pack its DNA. Epigenetics & Chromatin 9: 36.

81. Törönen P, Medlar A, Holm L. 2018. PANNZER2: a rapid functional annotation web server. Nucleic Acids Research 46: W84–W88.

82. Varley Á, Horkan HR, McMahon ET, Krasovec G, Frank U. 2023. Pluripotent, germ cell competent adult stem cells underlie cnidarian regenerative ability and clonal growth. Current Biology 33: 1–10.

83. Voigt O, Erpenbeck D, Worheide G. 2008. A fragmented metazoan organellar genome: the two mitochondrial chromosomes of Hydra magnipapillata. BMC Genomics 9: 350.

84. Wagner DE, Wang IE, Reddien PW. 2011. Clonogenic Neoblasts Are Pluripotent Adult Stem Cells That Underlie Planarian Regeneration. Science 332: 811–816.

85. Walker BJ, Abeel T, Shea T, Priest M, Abouelliel A, Sakthikumar S, Cuomo CA, Zeng Q, Wortman J, Young SK, et al. 2014. Pilon: An Integrated Tool for Comprehensive Microbial Variant Detection and Genome Assembly Improvement. PLOS ONE 9: e112963.

86. Wang D, King SM, Quill TA, Doolittle LK, Garbers DL. 2003. A new sperm-specific Na + /H + Exchanger required for sperm motility and fertility. Nature Cell Biology 5: 1117–1122.

87. Wang J, Sun H, Jiang M, Li J, Zhang P, Chen H, Mei Y, Fei L, Lai S, Han X, et al. 2021. Tracing cell-type evolution by cross-species comparison of cell atlases. Cell Reports 34: 108803.

88. Weismann A. 1883. Die Entstehung der Sexualzellen bei Hydromedusen (The origin of the sexual cells in hydromedusae). Gustav Fischer-Verlag, Jena.

89. Wessel GM. 2013. Getting schooled the ol’fashioned way. August Weismann and the “germ terms.” Molecular Reproduction and Development 80. https://onlinelibrary.wiley.com/doi/abs/10.1002/mrd.22149 (Accessed May 3, 2021).

90. Wheeler BM, Heimberg AM, Moy VN, Sperling EA, Holstein TW, Heber S, Peterson KJ. 2009. The deep evolution of metazoan microRNAs. Evolution & Development 11: 50–68.

91. Wong WY, Simakov O, Bridge DM, Cartwright P, Bellantuono AJ, Kuhn A, Holstein TW, David CN, Steele RE, Martínez DE. 2019. Expansion of a single transposable element family is associated with genome-size increase and radiation in the genus Hydra. Proceedings of the National Academy of Sciences 116: 22915–22917.

92. Young RA. 2011. Control of the Embryonic Stem Cell State. Cell 144: 940–954.

93. Zacharias H, Anokhin B, Khalturin K, Bosch TCG. 2004. Genome sizes and chromosomes in the basal metazoan Hydra. Zoology 107: 219–227.

94. Zimmermann B, Robb SMC, Genikhovich G, Fropf WJ, Weilguny L, He S, Chen S, Lovegrove-Walsh J, Hill EM, Chen C-Y, et al. 2022. Sea anemone genomes reveal ancestral metazoan chromosomal macrosynteny. 2020.10.30.359448. https://www.biorxiv.org/content/10.1101/2020.10.30.359448v2 (Accessed May 3, 2023).

